# Upper limb joint coordination acts to preserve hand kinematics after a traumatic brachial plexus injury

**DOI:** 10.1101/2022.09.06.506862

**Authors:** Luiggi Lustosa, Ana Elisa Lemos Silva, Raquel de Paula Carvalho, Claudia D. Vargas

## Abstract

**Background:** Traumatic brachial plexus injury (TBPI) causes a sensorimotor deficit in upper limb (UL) movements.

**Objective:** Our aim was to investigate the arm-forearm coordination of both the injured and uninjured UL of TBPI subjects.

**Methods:** TBPI participants (n=13) and controls (n=10) matched in age, gender, and anthropometric characteristics were recruited. Kinematics from the shoulder, elbow, wrist and index finger markers were collected while upstanding participants transported a cup to mouth and returned the UL to a starting position. The UL coordination was measured through the relative phase (RP) between arm and forearm phase angles and analyzed as a function of the hand kinematics.

**Results:** For all participants, the hand transport had a shorter time to peak velocity (p<0.01) compared to the return. Also, for the control and the uninjured TBPI UL, the RP showed a coordination pattern that favored forearm movements in the peak velocity of the transport phase (p<0.001). TBPI participants’ injured UL showed a longer movement duration in comparison to controls (p<0.05), but no differences in peak velocity, time to peak velocity and trajectory length, indicating preserved hand kinematics. The RP of the injured UL revealed altered coordination in favor of arm movements compared to controls and to the uninjured UL (p<0.001). Finally, TBPI participants’ uninjured UL showed altered control of arm and forearm phase angles during the deceleration of hand movements compared to controls (p<0.05).

**Conclusion:** These results suggest that UL coordination is reorganized after a TBPI so as to preserve hand kinematics.

## 1 Introduction

The brachial plexus consists in a dense network of spinal nerves originating from vertebrae C5 to T1. Traumatic brachial plexus injury (TBPI) occurs most commonly in young adults involved in motorcycle accidents (Faglioni et al., 2014), causing sensory, motor and autonomic deficits of the affected upper limb (Resnick, 1995). Brachial plexus nerve roots (C5-T1) can be partially or entirely affected (Dubuisson and Kline, 2002; Moran et al., 2005), with the degree of sensorimotor dysfunction varying as a function of the lesion extent and severity (Crouch et al., 2016). Proximal shoulder and elbow flexor muscles are the most susceptible to paralysis and sensory loss (Özkan and Aydin, 2001).

Although complete reconstruction of the damaged peripheral nerve pathways is not possible, complex reconstructive surgeries (Noland et al., 2019) and physical therapy (Kinlaw, 2005; Milicin and Sîrbu, 2018; Rich et al., 2019; Chagas et al., 2021) can help restore the motor function of the affected upper limb (UL). Usually, surgical procedures aim to recover shoulder abduction and external rotation, with greater focus on biceps strength restoration through nerve transfer (Hems, 2015). Surgery results show that most patients recover elbow flexion muscle strength to at least grade 3 (range: 0-5) in the Medical Research Council (MRC) (Sungpet et al., 2000; Teboul et al., 2004; Leechavengvongs et al., 2006). Restorative shoulder approaches, however, have a less successful result, and shoulder instability is observed in 50% of the patients after surgery (Hems, 2015).

Motion analysis has been used in the clinical context after a TBPI in order to quantify compensatory trunk movements and shoulder dysfunction, and thus help prioritize secondary surgical targets (Crouch et al., 2016; Webber et al., 2019; Nazarahari et al., 2020). For instance, Crouch et al. (2016) observed a reduced maximal strength for shoulder abduction and external rotation for injured upper limb movements, and Webber et al. (2019) identified limited external rotation of the shoulder when individuals with TBPI performed feeding and dressing tasks. Souza et al. (2021) analyzed the kinematic parameters of movement performed with the uninjured UL of individuals with TBPI in a free-endpoint whole-body reaching task requiring trunk motion. This task allowed the subjects to freely choose their final hand position, exposing them to a number of subjective choices (Haggard, 2008; Andersen and Cui, 2009; Berret et al., 2011; Hilt et al., 2016). Results revealed altered kinematic parameters for the uninjured UL of TBPI individuals when performing this task as compared to age-paired control participants.

TPBI was also shown to promote plastic modifications in the topographic organization of movement representations in the primary motor cortex (M1) (Mano et al., 1995; Iwase et al., 2001; Hsieh et al., 2002; Malessy et al., 2003; Pawela et al., 2010; Sokki et al., 2012; Yoshikawa et al., 2012; Liu et al., 2013; Qiu et al., 2014; Fraiman et al., 2016; Bhat et al., 2017). Functional Magnetic Resonance Imaging (fMRI) in resting state of TPBI individuals showed reduced interhemispheric connectivity in M1 (Liu et al., 2013), reduction in the connectivity of arm and hand representations of M1 with the ipsilateral supplementary motor area (Qiu et al., 2014), and reduced local connectivity of the UL and trunk representations in M1 at both hemispheres (Fraiman et al., 2016). Since studies with animals have already shown that the different UL representations (shoulder, elbow, wrist and fingers) have a strong overlap and mingled distribution (Kwan et al., 1978; Huntley and Jones, 1991; Park et al., 2001), these alterations in the connectivity of M1 could have an effect on motor planning. Accordingly, Rangel et al. (2021) showed that the EEG activity associated with predicting an upcoming event was altered bilaterally in the sensorimotor cortex of TBPI individuals. Taken together, these results point towards plastic modifications of UL motor plans after a TBPI.

Motor plans of goal-directed actions have been classically accessed through kinematic measurement (Bernstein, 1967; Soechting and Lacquaniti, 1981; Marteniuk et al., 1987; Papaxanthis et al., 1998; Desmurget et al., 1999; Svoboda and Li, 2018). The motor plan is thought to encode both where the reach will land on average (the endpoint) and the expected movement duration (Wolpert and Landy, 2012). Reaching movements have been shown to display regularities such as typical straight trajectories and bell-shaped velocity profiles (Bernstein, 1967; Atkeson and Hollerbach, 1985; Flash and Hogan, 1985; Marteniuk et al., 1987; Soechting and Flanders, 1991). Duration is an important kinematic component of such motor decisions because of the speed-accuracy tradeoff (Wolpert and Landy, 2012). In addition, Marteniuk et al. (1987) showed that when the task demands greater precision, the duration of the deceleration phase of the trajectory is increased as a consequence of the greater demand for sensory feedback to perform the task.

Different studies have shown that this straight trajectory and smooth control of hand velocity is made by the coupling of shoulder and elbow joint movements (Morasso, 1981; Soechting and Lacquaniti, 1981; Atkeson and Hollerbach, 1985). This means that the motor systems must coordinate the muscles acting at shoulder and elbow joints to produce a controlled rotation of the arm and the forearm segments, resulting in a slightly invariant hand trajectory (Soechting and Lacquaniti, 1981). This inter-joint coordination has been assessed in UL kinematic analysis by different methods, such as measuring a correlation coefficient between joints’ angular displacement (Murphy et al., 2006, 2011; de los Reyes-Guzmán et al., 2014) or the ratio between joints’ range of motion (Bagesteiro et al., 2020). However, that type of measure does not allow a temporal analysis of the pattern of coordination during a task. A more detailed analysis of these patterns can be made through the relative phase, which is a dynamic systems approach that considers two anatomically linked body segments as a coupled system acting to move an effector efficiently (Kelso, 1995; Barela et al., 2000; Lamb and Stöckl, 2014). The measure compresses the displacement and velocity of two different segments in a single variable yielding a measure that shows how fast a segment is moving in relation to the other for every instant of the movement (Clark and Phillips, 1993; Barela et al., 2000). The coordination of arm-forearm kinematics has already been described employing the relative phase parameter in the context of joint angle variability after UL fatigue (Yang et al., 2018), clinical assessment of motor coordination of the affected UL in stroke patients (Daunoravičienė et al., 2017), analysis of bimanual interlimb coordination (Liddy et al., 2017) and interlimb coordination in sports performance (Guignard et al., 2017). As a useful tool in describing UL kinematics, the relative phase could be important in clarifying the strategies of motor control after a TBPI.

An impairment of shoulder and elbow muscles could change the UL pattern of coordination of TBPI individuals. The most usual form of TBPI spares hand movements but leaves a strength deficit in shoulder and elbow muscles even after surgeries (Hems, 2015). This weakness in the muscles could create new constraints to joint motion, requiring an adaptation of previous learned coordination patterns for hand control. Adaptations of this type have been shown in animal models after peripheral nerve injury (Chang et al., 2009, 2018; Sabatier et al., 2011; Bauman and Chang, 2013). It has been observed that recovery promotes a new combination of joint angles of the affected paw in an attempt to preserve limb function in gait, suggesting that joint coordination is reorganized in order to conserve effector performance (Chang et al., 2009, 2018; Sabatier et al., 2011; Bauman and Chang, 2013). Likewise, the movement of injured UL in TBPI individuals could show a modified pattern of arm-forearm coordination combined with preservation of hand kinematic performance. Furthermore, as changes in the kinematics of the uninjured UL in TBPI have been previously reported (Souza et al., 2021), we also investigated the arm-forearm coordination and its relationship with the hand kinematics of this limb.

The main objective of this study was to analyze the coordination pattern of the UL in TBPI individuals, compared with control individuals without TBPI while they performed the movement of bringing a cup to the mouth (transport and return). In a regular reaching movement, shoulder muscles must activate in advance to stabilize arm and forearm motion (Ricci et al., 2015). However, the weakness in shoulder muscles after a TBPI could make it difficult to stabilize UL motion. In more unstable distal movements, healthy subjects reduce distal joint motion and increase proximal joint motions (van der Kamp and Steenbergen, 1999). Accordingly, we conjectured that the pattern of arm-forearm coordination of the injured UL would be modified so as to preserve the hand kinematic performance. More specifically, the analysis of the arm-forearm coordination could reveal if the UL movements rely more on the arm as compared to the forearm segment both for the injured and the uninjured UL as a result of TBPI-induced plastic modifications in motor plans (Souza et al., 2021; Rangel et al., 2021).

## 2 Materials and methods

### 2.1 Participants

From June 2018 to August 2020, TBPI patients from a database maintained by the Laboratory of Neuroscience and Rehabilitation of the Federal University of Rio de Janeiro were invited to participate in the study. This database contains epidemiological, physical, clinical and surgical information of a large cohort of TBPI patients (Patroclo et al., 2019). The following inclusion criteria were used to select the patients to participate in this study: unilateral TBPI diagnosed by clinical evaluation or complementary exams, age between 18 and 60 years, right-hand dominance before TBPI verified with Edinburgh Handedness Inventory (Oldfield, 1971). Exclusion criteria were: obstetrical brachial plexus injury, visual loss or uncorrected deficits, and presence of neurological diseases. The functionality of the upper limb was measured with the Brazilian Portuguese version of Disabilities of the Arm, Shoulder and Hand questionnaire (DASH) (Orfale et al., 2005). This questionnaire is composed of 30 questions addressing the ability to perform daily activities with the UL and the severity of symptoms. The final score is within a range from 0 to 100. The higher the score, the greater the disability.

Controls matching in age, height and weight, and without any report of musculoskeletal or neurological problems were recruited to compose the control group. Participants were verbally informed about all experimental proceedings and signed a written consent to join the tests. The ethics committee of Institute of Neurology Deolindo Couto at the Federal University of Rio de Janeiro approved all experimental procedures (Plataforma Brasil, CAEE: 51657615.6.0000.5261; process number: 1.375.645).

### 2.2 Kinematic recording of the task

Seven motion capture cameras with 1.0 megapixel resolution (Vicon Bonita 10, Vicon, USA) and the Vicon Nexus software version 2.2 (Vicon Motion Systems Ltd, Vicon, USA) were used to collect three-dimensional movements at the sampling rate of 100 Hz. Four reflective 15 mm markers were placed on the following structures of participants’ UL: apex of index finger, ulna styloid process, lateral epicondyle of the humerus and acromion (Figure 1B).

**Figure 1.**
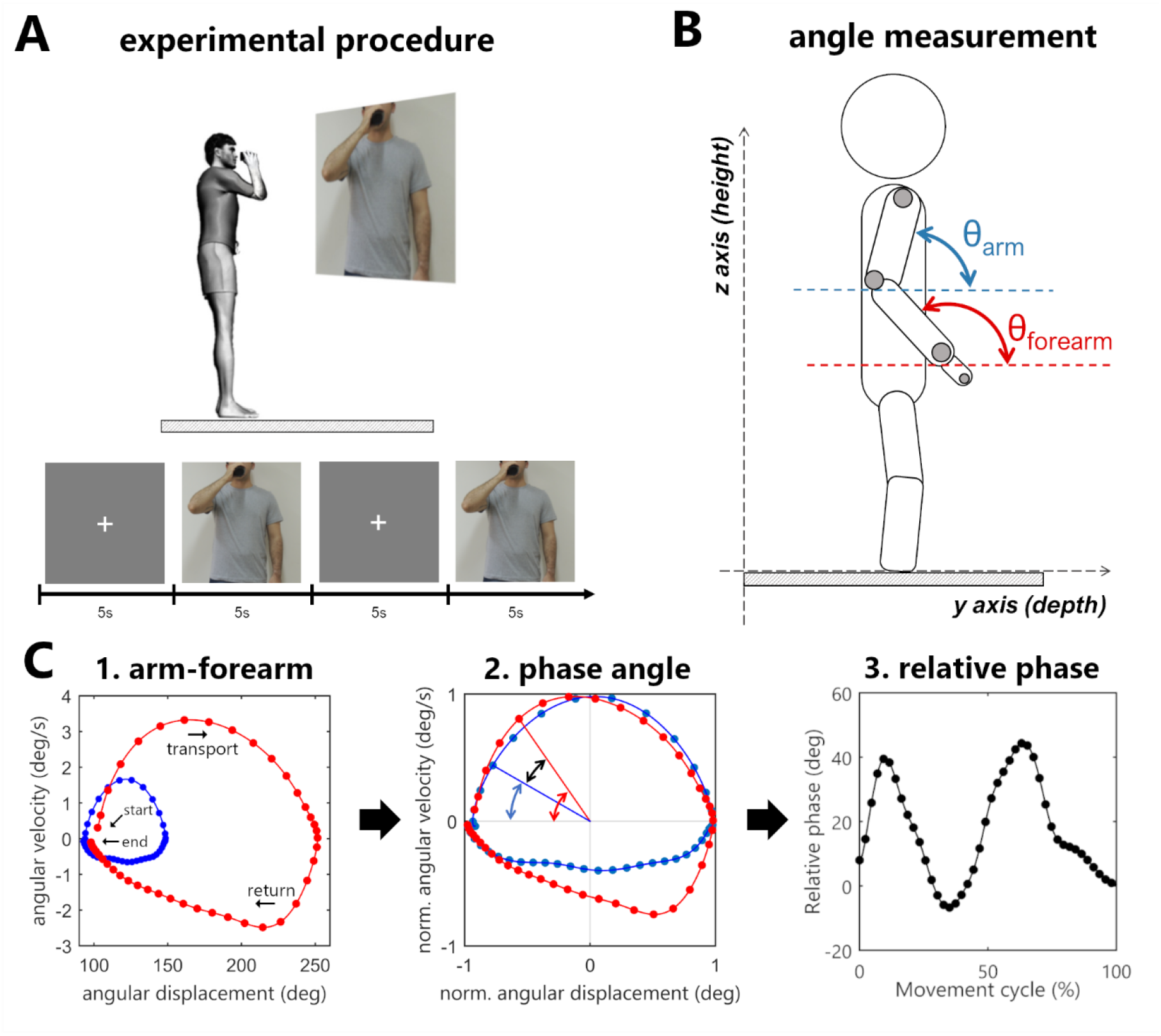
**(A)** Experimental procedure. The sequence of images consisted of a fixation cross and a picture of a person holding a cup in front of the mouth. When the movement image was displayed, participants had to immediately bring the cup to the mouth and then return to the standby position. **(B)** Angle measurement in sagittal plane. **(C)** Arm-forearm dynamics analyzed during the cycle of transporting the cup to the mouth and returning to the standby position (blue points: arm, red points: forearm). **(C1)** Starting and ending points of the full cycle are signalized by arrows. The evolution is seen in a clockwise direction. The points in the graph represent the angular displacement (θ) and the angular velocity (ω) of a segment in a time instant. **(C2)** The difference between the normalized phase angles (black arrow) is the relative phase (RP = ФForearm - ФArm). **(C3)** The relative phase over time.

Participants stood up over a rigid surface placed in the motion capture area with their feet positioned hip-width apart and parallel to the sagittal plane. They were asked to hold an empty plastic cup in the tested UL hand. The cup was rigid (not deformable) and weighted 200 g (dimensions: length: 13 cm, smaller diameter: 5 cm, larger diameter: 7 cm). The Presentation software (Neurobehavioral System, Inc., USA) was used to project (Epson PowerLit S18+ ®, Epson, Japan) two subsequent pictures to the participants: a fixation cross followed by a picture of a person holding a plastic cup in front of his mouth (Figure 1A).

Participants were instructed to hold the plastic cup and keep an upright position with upper limbs relaxed by their sides every time the fixation cross was shown. When the movement figure was displayed, they were asked to reproduce the end position exhibited in the figure by bringing the plastic cup to the mouth at a comfortable self-selected speed and immediately returning to the stand-by posture (i.e., they should not wait for the fixation cross to return) (Figure 1A). Each figure was displayed to the participants for five seconds, during which the recording of the participants’ UL movements was performed.

Before the kinematic recording, two trials were performed as training and the data was not included in the analysis. The task was executed in two blocks of eight trials per UL. For control participants, the first block was applied to the right UL, and the second block was applied to the left UL. For TBPI participants the first block was applied to the uninjured UL and the second block to the injured UL. To perform the trials with the injured side, TBPI participants had to score at least 3 in the MRC scale for the elbow flexor muscles and the hand had to be strong enough to hold the cup firmly. The muscle strength of other UL muscle groups was obtained for all participants (except P14 and P15) from a follow up assessment performed by the laboratory staff.

### 2.3 Data analysis

An offline processing was done in the Vicon Nexus 2.2 software for the reconstruction of reflective marker coordinates in three-dimensional space (mediolateral, X, antero-posterior, Y and vertical, Z) and for correction of gaps in the capture process. Processed data was exported to MATLAB software (R2015a, Mathworks Inc., Natick, MA, USA), and a 5th-order low pass filter at 10Hz cutoff was applied to marker data before calculating desired variables. The task was divided in two phases: the transport phase of the plastic cup from standby position to the mouth, and the return phase to standby position. As the index finger displayed a curved trajectory, its speed was estimated by its tangential velocity 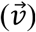 in relation to the path. For each instant of time, the tangential velocity 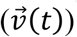 was calculated by multiplying the sampling frequency (*f*_*s*_) to the magnitude of the index finger displacement vector 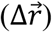, which corresponds to the index finger displacement in 3-D space.

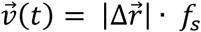

The beginning of a movement phase was determined as the moment at which the tangential velocity of the index finger marker exceeded 5 % of peak velocity, and the phase ending as the moment at which the velocity dropped below 5% of peak velocity (Esteves et al., 2016). After the phase detection, the velocity was time-normalized by a linear interpolation of 400 points. An inspection of velocity profiles was executed to exclude trials suggesting evident processing errors or trials in which participants did not complete the full movement within the expected time (up to 5 sec). All trials that matched these criteria were excluded before statistical analysis.

#### 2.3.1 Hand kinematics outcomes

Hand kinematic performance was estimated based on index finger movements (see Souza et al., 2021 for more details). Movement duration (MD) was determined as the time elapsed between the beginning and the end of a movement phase. Peak velocity (PV) was the maximum index finger velocity in a movement phase. Time to peak velocity (TPV) was calculated as the ratio between the time spent to reach peak velocity and the movement duration. This ratio represents the percentage of MD spent in hand acceleration. Trajectory length (TL) was measured as the index finger traveled distance in a movement phase.

The number of movement units (NMU) is a local maximum in the index finger velocity curve. A movement unit was set when the difference between a minimum value and its next maximum exceeded 20 mm/s (Bustrén et al., 2017). Usually, hand reaching movements have only a single peak velocity, and a greater number of peaks indicates a loss in smoothness (de los Reyes-Guzmán et al., 2014). Normalized end height (NEH) was calculated as the ratio between the index finger height at the end of movement and the participant’s height. This variable was calculated only for the transport phase.

#### 2.3.2 Arm - forearm coordination outcomes

Because the focus of our study was to analyze the arm-forearm coordination, we devised the task of bringing a cup to the mouth, expecting that TBPI participants would succeed in performing the movement despite their limitations.

The arm-forearm coordination was assessed through phase angles that assume a fixed two-dimensional plane (Barela et al., 2000). In the cup to mouth task, which describes a cyclic movement, most of the motion of the arm and forearm segments occurs in the sagittal plane, so in order to assess arm-forearm coordination changes after a TBPI we selected this plane to analyze the phase angles.

In this analysis, the angular position of a segment in space is plotted against its angular velocity, thus every point of this graph represents the displacement and velocity of a given segment (Figure 1C). For this purpose, vector for the segment (arm or forearm) was calculated by subtracting the coordinates of two markers in the YZ plane. The arm vector was calculated using the humerus lateral epicondyle and acromion markers, and the forearm with styloid process of ulna and humerus lateral epicondyle markers. The angular displacement of the segment in relation to the YZ plane (θ) was calculated using the arc tangent of two arguments, and the segmental angular velocity (ω) was obtained by the first derivative of angular displacement.

Next, the angular displacement (θ) and the angular velocity (ω) were normalized, limiting the range of the signal between -1 and 1 (Barela et al., 2000; Lamb and Stöckl, 2014). The angular displacement normalization uses its maximum and minimum values as reference, thus after the transformation, the zero value represents the midway between the greatest and lowest value in the signal. The angular velocity is normalized based on its maximum, allowing the zero value to have the same meaning it had prior to the normalization process.

Finally, using normalized angular displacement and normalized angular velocity as arguments of the arc tangent function, a polar angle was obtained. The subtraction of this value by 180° resulted in the phase angle of the segment (Ф_Forearm_, Ф_Arm_). The arm-forearm relative phase (RP) was then calculated by the subtraction of the phase angles (RP = Ф_Forearm_ - Ф_Arm_). A positive RP indicates that the forearm has a greater displacement and velocity in relation to the arm, and the inverse when the RP is negative. Raises in the RP will always come from the increased difference between phase angle of the arm and forearm, indicating an increase in the contribution of the forearm to the movement. Likewise, reductions in the relative phase will indicate an increase in the contribution of the arm to the movement. These three continuous measures of segmental dynamics (Ф_Arm_, Ф_Forearm_, and RP) were time-normalized by a linear interpolation of 400 points for the comparison between groups.

Phase angles (Ф_Forearm_, Ф_Arm_) were plotted along time, to analyze the arm-forearm controlling strategy. The curves of the segments phase angles over time were splitted in two parts according to hand acceleration and deceleration, and next, the area under these curves was calculated using their absolute values. A measurement was designed specifically to the RP. Ten percent (10%) of signal samples were collected in three different moments (time windows) of the hand kinematics (movement start, hand peak velocity, and movement ending) to search for differences in the coordination pattern within the movement cycle.

### 2.4 Statistical analysis

Statistical analysis was conducted using Jamovi 1.6.23 (The Jamovi Project) and GraphPad Prism 7 (San Diego, California, USA). Normal distribution was tested through Shapiro Wilk tests, and non-parametric tests were performed when necessary. The level of significance of the study was set at *p* < 0.05. The eight trials performed by each UL were averaged and a final mean and standard deviation (SD) was calculated for each group. TBPI UL analysis was divided into two groups: uninjured UL and injured UL. Left and right UL performance in controls was compared to check the presence of an effect for the movement side. No significant differences (Mann-Whitney U test *p* > 0.05) between left and right sides were observed for any of the outcome variables. Therefore, the performance of control individuals was calculated as the average between their right and left UL. Since the injury in the right UL could have an impact on the handedness of participants with right side TPBI, we compared the UL kinematic performance of participants according to their side of TPBI (left or right). No significant differences (Mann-Whitney U test p > 0.05) between left and right sides of TPBI were observed for any of the outcome variables, both for the injured and uninjured UL.

A two-way ANOVA was applied to hand kinematic variables (MD, PV, TPV, TL) with the movement phase (transport or return) and group (control, uninjured UL, and injured UL) as factors. Tukey test was applied for *post hoc* multiple comparisons. A one-way ANOVA was applied to NEH, because this variable was only measured for the transport phase. To compare the variances between groups the Bartlett test was used.

For the comparison of the area under phase angle curves, the movement phase (transport or return) was not taken as a factor because the area increases constantly during the movement cycle. Then, a one-way ANOVA was conducted to compare the groups. Tukey test was used as a *post hoc* for multiple comparisons. For non-normal distribution, Kruskall-Wallis test was used and Dwass-steel-critchlow-fligner pairwise comparisons as *post hoc*. A three-way ANOVA was used in relative phase statistics with the movement phase (transport or return), hand kinematic moments (start, peak velocity, end) and group as factors. Tukey test was applied as a *post hoc*.

## 3 Results

### 3.1 Participants

Thirteen patients (n=13) with TPBI matched inclusion criteria and were selected to the study. Their median age was 35 years (range: 20-55 years), median weight 85.00 kg (range: 39.00-105.00 kg), median height 1.74 m (range: 1.52-1.84 m). Ten control (n=10) participants matched with TBPI participants were selected for the study. Their median age was 27.5 years (range:19-58 years), median weight 78.45 kg (range: 52.00-104.20 kg), median height 1.78 m (range:1.53-1.90 m). No significant differences in age, weight, and height were found between TBPI participants (n=13) and controls (Mann Whitney U test *p* > 0.05 for all variables). TBPI data concerning age, injury, surgery and rehabilitation status are summarized in Table 1. From the thirteen TBPI participants, seven were injured on the right side and six on the left side. Six TBPI individuals were injured at upper trunk level extending to C7 (C5, C6 and C7 nerve roots), four individuals had a total plexus injury (nerve roots from C5 to T1), one individual had an upper trunk injury level (C5-C6 nerve roots), and one had a posterior cord injury involving the axillary nerve. The median time elapsed from injury was: 3.42 years, range: from 2 months to 9 years and 8 months. Eleven TBPI individuals underwent surgical procedures (median time elapsed from surgery: 3.50 years, range: from 6 months to 8 years and 5 months). Muscle strength of the injured UL in TBPI individuals measured with the MRC scale is summarized in Table 2. The variability observed in UL strength of TBPI participants goes along with the heterogeneity observed in the extent of the injury and type of surgical procedure. From the thirteen TBPI participants, six (n=6) scored at least 3 out of 5 for elbow flexors (see Table 2) in the MRC scale and were able to perform the trials with the injured UL. Two TBPI participants were not enrolled in any rehabilitation program. No significant differences in age, weight, and height were found between TBPI participants in the injured UL group (n=6) and controls (Mann Whitney U test p > 0.05).

**Table 1.** TBPI Individual Characteristics

| ID | Age | Injury Level | Injury Side | Time from injury | Surgery | Time from surgery | DASH score | Rehab |
| --- | --- | --- | --- | --- | --- | --- | --- | --- |
| P02 | 37 | C5-C7 | L | 1y, 4m | Ac-SE<br>Oberlin | 6m | 50.8 | Y |
| P03* | 42 | C5-C7 | L | 3y | Oberlin | 2y, 6m | 27.5 | Y |
| P04 | 29 | C5-T1 | L | 3y, 5m | INT-MSC<br>Ac-SE | 3y, 2m | 43.3 | Y |
| P05* | 28 | C5-C7 | R | 4y, 3m | Ac-SE<br>Oberlin | 3y, 10m | 59.5 | Y |
| P06 | 29 | C5-T1 | R | 4y, 6m | INT-MSC<br>Ac-SE | 3y, 4m | 30.0 | Y |
| P07 | 39 | C5-C7 | L | 2y, 9m | Not specified<sup>1</sup> | – | 20.0 | Y |
| P08 | 27 | C5-C7 | R | 2y, 2m | Ac-SE | 11m | – | N |
| P09 | 38 | C5-T1 | R | 9y, 8m | INT-MSC<br>Ac-SE | 8y, 5m | 59.2 | Y |
| P11* | 30 | C5-C6 | R | 4y, 4m | Oberlin<br>Tr-AX<br>Ph-SE | 3y, 11m | 45.0 | Y |
| P12 | 35 | C5-T1 | R | 4y | Ac-MC | 3y, 6m | 45.7 | Y |
| P13* | 43 | C5-C7 | L | 5y, 7m | Ac-SE<br>Oberlin | 5y, 2m | 25.0 | N |
| P14* | 20 | PC+Ax | R | 0y, 4m | None | – | – | Y |
| P15* | 55 | C5-C6 | L | 0y, 2m | None | – | – | Y |

\* TBPI participants who were able to perform the task with the injured UL. Age in years; Injury level (PC+Ax.: posterior cord injury with neurotmesis of axillary nerve); Upper limb side of injury (L: left; R: right); Time elapsed from injury to experiment date; Underwent surgeries (Oberlin: ulnar nerve transfer to musculocutaneous nerve; INT-MSC: intercostal nerve transfer to musculocutaneous nerve; Ac-SE: accessory nerve transfer to suprascapular nerve; Ph-SE: phrenic nerve transfer to suprascapular nerve; Tr-Ax: medial triceps transfer to axillary nerve; Ac-MSC: accessory nerve transfer to musculocutaneous nerve; <sup>1</sup>Not specified: surgery not specified in medical records). Time elapsed from the last surgery to the experiment date; DASH score (Disabilities of the Arm, Shoulder and Hand questionnaire); Enrollment in rehabilitation (Y: yes, N:no);

**Table 2.** TBPI patients’ evaluation of the injured UL muscle strength

| ID | Shoulder flexor | Shoulder extensor | Shoulder external rotator | Elbow flexors | Elbow extensors | Wrist extensors | Finger flexors | Finger abductors |
| --- | --- | --- | --- | --- | --- | --- | --- | --- |
| P02 | 0 | 0 | 0 | 0 | 0 | 3 | NT | NT |
| P03* | 4 | 4 | 4 | 4 | 4 | 2 | 4 | 2 |
| P04 | 0 | 5 | 0 | 0 | 0 | 0 | 0 | 0 |
| P05* | 2 | 5 | 0 | 3 | 5 | 5 | 5 | 5 |
| P06 | 0 | 5 | 2 | 2 | 0 | 0 | 0 | 0 |
| P07 | 0 | 5 | 2 | 2 | 4 | 5 | 5 | 5 |
| P08 | 0 | 5 | 0 | 2 | 3 | 0 | 5 | 3 |
| P09 | 0 | 5 | 0 | 2 | 2 | 2 | 5 | 2 |
| P11* | 4 | 5 | 5 | 5 | 5 | 5 | 5 | 5 |
| P12 | 2 | 5 | 1 | 5 | 0 | 0 | 0 | 0 |
| P13* | 0 | 5 | 1 | 5 | 4 | 5 | 5 | 3 |
| P14* | NT | NT | NT | 3 | NT | NT | NT | NT |
| P15* | NT | NT | NT | 3 | NT | NT | NT | NT |

Muscular Manual Test (Medical Research Council): 0 - No visible or palpable contraction; 1 - Visible or palpable contraction with no motion; 2 - Full range of movement with gravity eliminated; 3 - Full range of movement against gravity; 4 - Full range of movement against gravity, moderate resistance; 5 - Full range of movement against gravity, maximum resistance. NT – not tested. \* TBPI participants who were able to perform the task with the injured UL. P14 and P15 had only their elbow flexors evaluated.

### 3.2 Hand kinematic performance

A two-way ANOVA taking groups (controls, injured UL and uninjured UL) and movement phase (transport or return) as factors showed group differences for MD (F_(2,52)_ = 3.62, *p* < 0.05). *Post hoc* analysis showed that MD was longer for the injured UL (*p* < 0.05) when compared to controls. Moreover, a main effect for the movement phase was observed in TPV (F_(1,52)_ = 7.99, *p* < 0.01). *Post hoc* analysis showed that subjects reached the PV earlier in hand transport when compared to the return movement (*p* < 0.01). This lower TPV indicates a prolonged deceleration movement (Figure 2). The TL also presented a main effect for movement phase (F_(1,52)_ = 11.34, *p* < 0.001), and *post hoc* revealed that the TL was shorter for the transport phase in comparison to the return (*p* < 0.001). No significant differences were observed in the PV for group (F_(2,52)_ = 2.67, *p* > 0.05) or phase (F_(1,52)_ = 3.05, *p* > 0.05). Calculated means and SD for each group are reported in the Supplementary Table S1.

**Figure 2.**
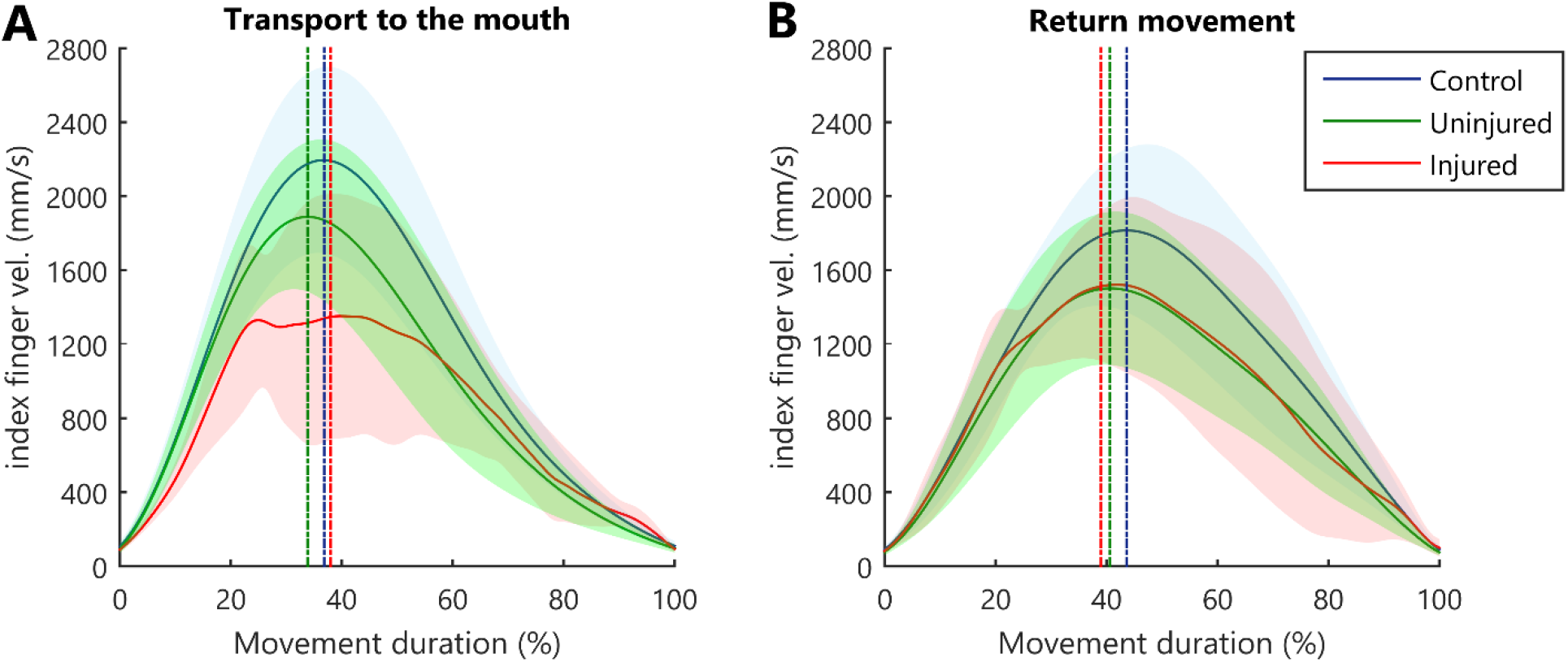
Kinematic profile of hand movement. Average index finger velocity (solid line), and SD (shaded area). The straight vertical dashed line indicates the mean instant for hand PV.

The SD seen in the velocity curve suggested that the injured UL (Figure 2, red shaded areas) showed a more diversified pattern of velocity profiles when performing movements. Therefore, we compared the variance between groups in the different movement phases. During the transport to the mouth, significant differences between groups were observed in the TPV (B_(2)_ = 15.65, *p* < 0.001, B = 15.65 > Χ^2^ (5.99)), but not in MD (B_(2)_ = 4.50, *p* = 0.106, B = 4.50 < Χ^2^ (5.99)), PV (B_(2)_ = 2.01, *p* = 0.367, B = 2.01 < Χ^2^ (5.99)) or TL (B_(2)_ = 1.45, *p* = 0.484, B = 1.45< Χ^2^ (5.99)). In the return of the hand to the starting position, no significant differences were observed between groups for MD (B_(2)_ = 0.59, *p* = 0.743, B = 0.59 < Χ^2^ (5.99)), PV (B_(2)_ = 0.49, *p* = 0.782, B = 0.49 < Χ^2^ (5.99)), TPV (B_(2)_ = 5.67, *p* = 0.059, B = 5.67 < Χ^2^ (5.99)) and TL (B_(2)_ = 1.73, *p* = 0.421, B = 1.73 < Χ^2^ (5.99)).

The hand normalized end height (NEH) differed significantly between groups (F_(2,26)_ = 6.97, *p* < 0.05). *Post hoc* analysis indicated that injured UL movements ended in a lower NEH compared to controls (*p* < 0.05) and to the uninjured UL (*p* < 0.01) (Supplementary Table S1). In both phases of movement (transport and return), all participants of the control and the uninjured UL group exhibited only one peak in velocity signal (NMU=1). The injured UL exhibited a higher NMU both in transport (mean: 1.73, SD: 1.70) and return (mean: 1.54, SD: 1.02). No statistical test was performed to this variable because all participants in the control and uninjured UL group had only a single peak velocity (NMU=1).

### 3.3 Arm and forearm phase angles

Phase plots for the arm and forearm segments can be observed in Figure 3A and 3B, respectively. During the transport phase acceleration, there was a significant difference in the arm phase angle area between groups (F_(2,26)_ = 26.52, *p* < 0.001). *Post hoc* tests indicated that the injured UL had a greater area in comparison to controls (*p* < 0.001) and to the uninjured UL (*p* < 0.001) (Figure 3C). In the deceleration of the transport phase, there was a difference for forearm phase angle area (H_(2)_ = 6.59, *p* < 0.05) (Figure 3D), and multi-comparisons test revealed a greater area for uninjured limb in comparison to controls (*p* < 0.05). No significant differences were observed in the arm phase angle area during cup transport deceleration (H_(2)_ = 0.39, *p* = 0.821) (Figure 3C), and in the forearm phase angle area during hand acceleration (F_(2,26)_ = 1.09, *p* = 0.35) (Figure 3D).

**Figure 3.**
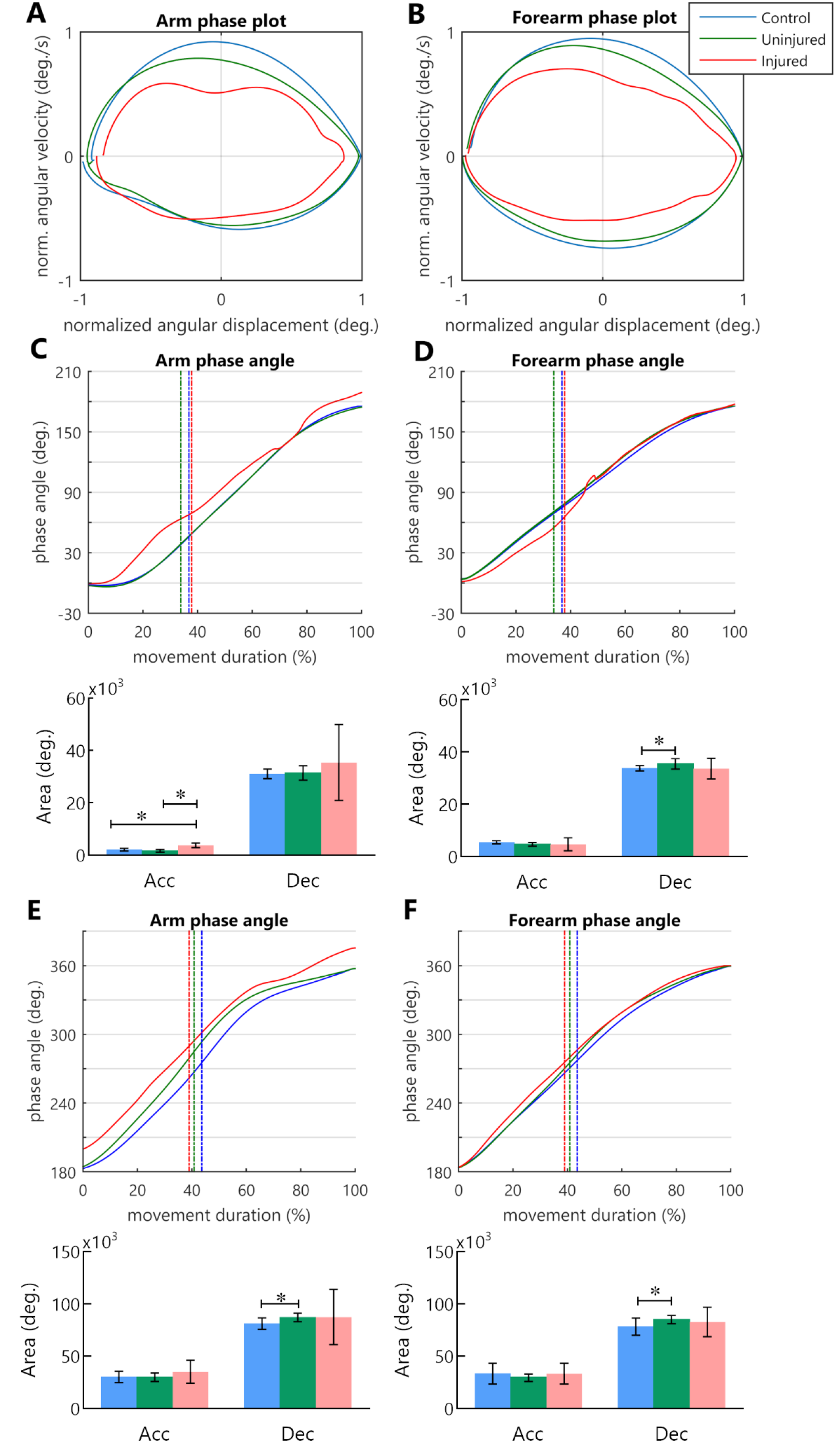
Phase angle of arm and forearm kinematics. **(A**,**B)** Arm and forearm phase plots. **(C**,**D**,**E**,**F)** Arm and forearm dynamics during the cycle of movement. Mean phase angle (filled line) is plotted for each group and the vertical dashed line indicates the moment in which the hand achieves the peak velocity. The result of area calculation is expressed in a bar graph under each phase angle plot. **(C**,**D)** Arm and forearm phase angle during transport to mouth. **(E**,**F)** Arm and forearm phase angle in the return to the starting position.

When the UL returned to the standby position, differences in the phase angle areas were observed solely during the deceleration of the hand. There was a difference between groups for arm phase angle area (H_(2)_ = 8.99, *p* < 0.05) and forearm phase angle area (H_(2)_ = 6.51, *p* < 0.05). In both segments, *post hoc* multiple comparisons revealed that the uninjured UL area was significantly larger than that of controls (*p* < 0.05) (Figure 3E, 3F). During hand acceleration, no differences between groups were found for arm phase angle area (H_(2)_ = 2.06, *p* = 0.357) and forearm phase angle area (H_(2)_ = 5.84, *p* = 0.054).

### 3.4 Relative phase in arm-forearm coordination

The RP measure along the movement cycle showed a specific coordination pattern across control and uninjured UL groups during the transport to mouth (Figure 4A, 4B). When the hand began to move towards the mouth the RP became progressively more positive until moments before the peak velocity. This positive RP represents a higher forearm displacement and velocity in relation to the arm during almost the entire hand acceleration. Near the peak velocity, there was a shift in the RP, with a continuous decrease until the end of the transport phase (Figure 4A, 4B), indicating that there was a decrease in forearm predominance during hand deceleration. In contrast, diversified patterns of coordination were found for the injured UL movement among TBPI participants, resulting in high SD values for the RP in this group (Figure 4C).

**Figure 4.**
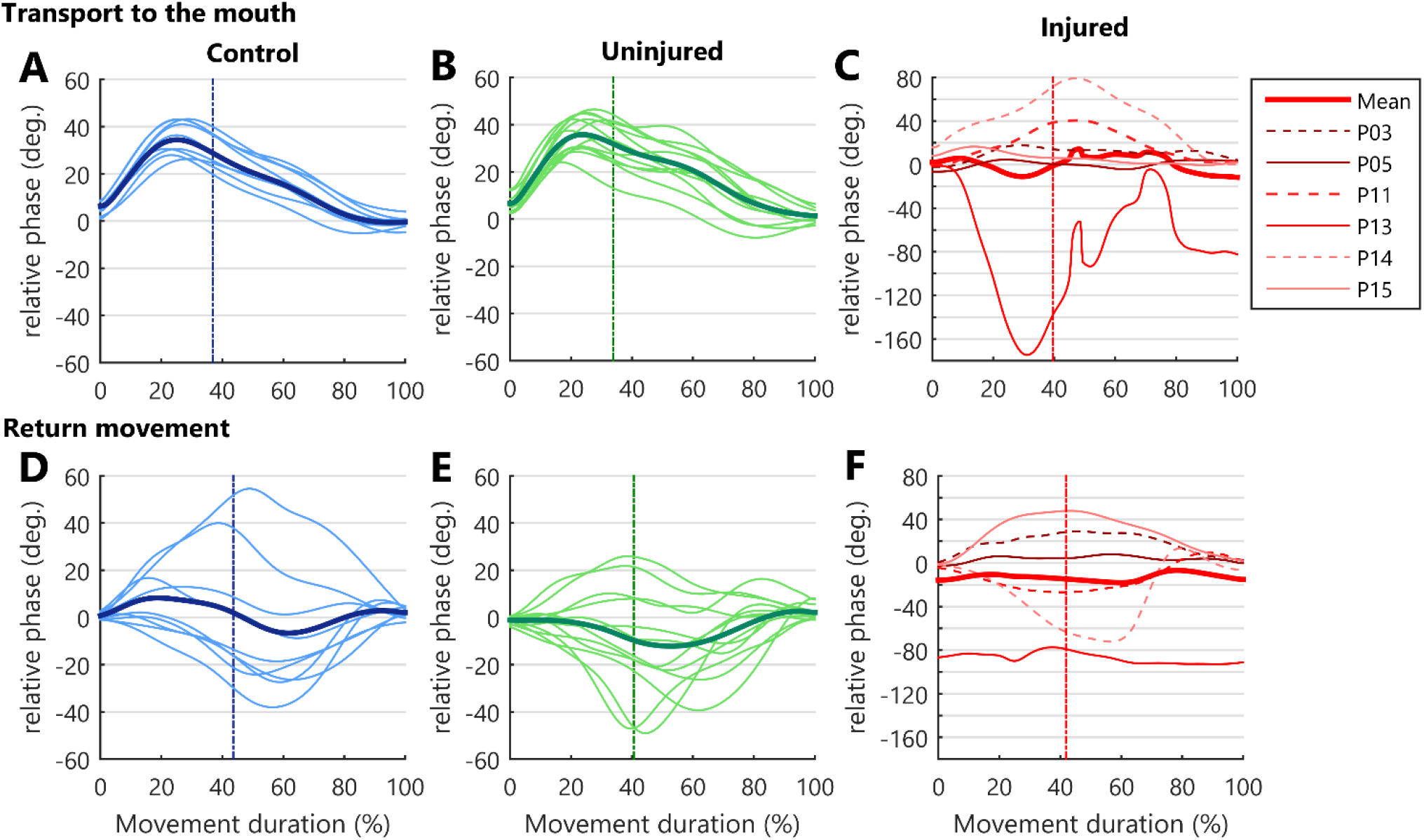
Upper limb intersegmental coordination. RP curves during the bringing a cup to mouth cycle (transport and return) and expressed as a function of the percent of movement phase duration. Solid thin lines correspond to the mean RP of each participant and the bold thick line correspond to the mean performance of the group. The vertical dotted line indicates the moment when the hand achieves the peak velocity. As the injured upper limb exhibited a higher RP variability, a grid line was marked in Y axis for each 20 degrees. **(A**,**B**,**C)** RP in hand transport to the mouth.. **(D**,**E**,**F)** Relative phase in the return to starting position.

During the return to the standby position, more diversified patterns of coordination were observed in all groups, contrasting to the transport to mouth phase. This behavior was seen as a larger SD in groups RP (Figure 4D, 4E, 4F).

Discretized RP measures were submitted to a three-way ANOVA considering groups, moment of hand kinematics (start, peak velocity, end), and movement phase (transport or return) as factors (Supplementary Table S2). Results indicated an effect for group (F_(2,156)_= 7.43, *p* < 0.001), and *post hoc* showed a more negative RP for the injured UL when compared to controls (*p* < 0.001) and to the uninjured UL (*p* < 0.05), pointing to a greater arm use in injured UL movements (Figure 5A).

**Figure 5.**
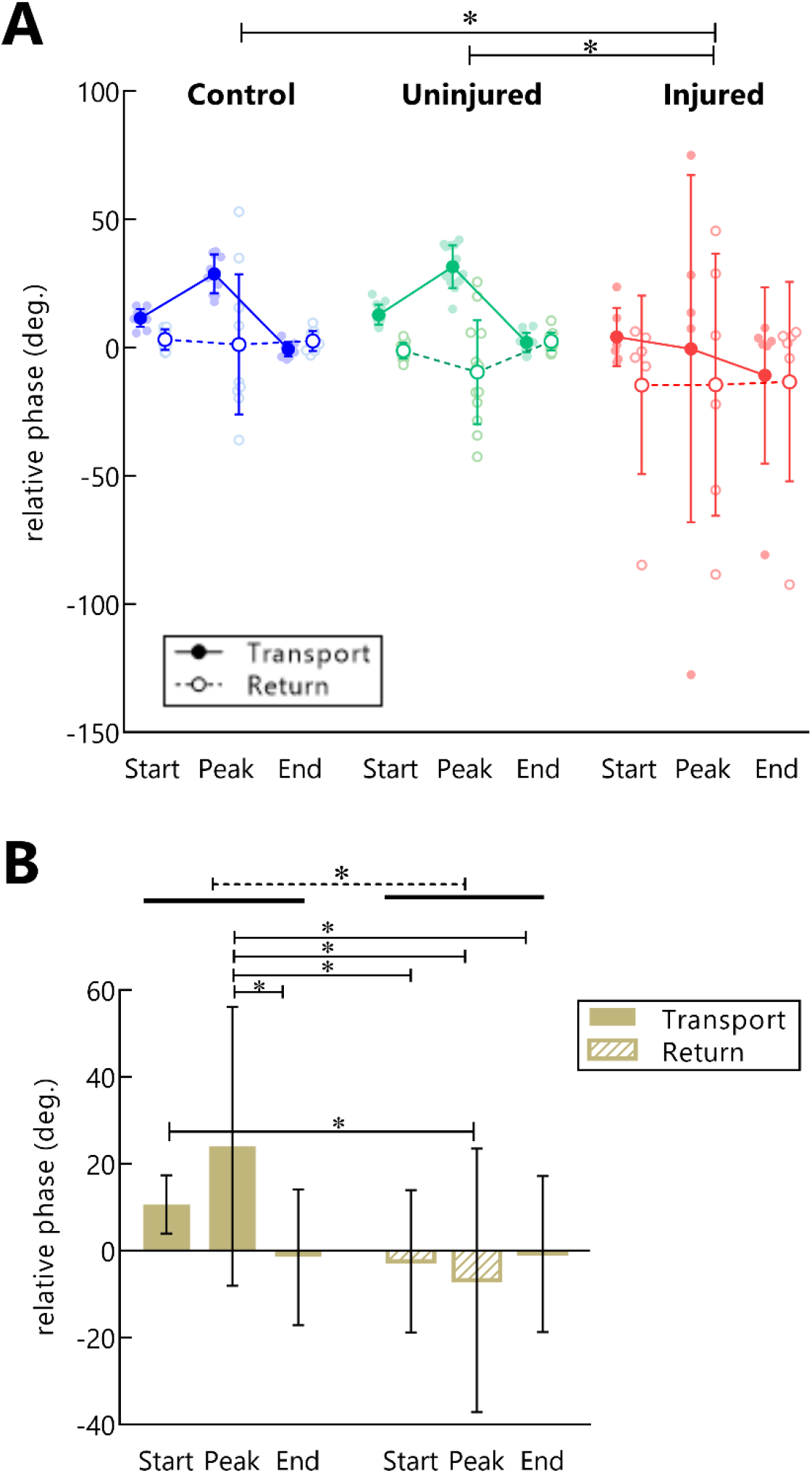
RP in control, injured and uninjured UL groups according to movement (transport or return) and instants of hand movement (start, peak velocity, end). **(A)** Each point represents the average RP per subject in start, peak velocity, end of movement in hand transport to the mouth (filled dots) and return to the starting position (open dots) **(B)** Main effect for movement phase. Mean RP is expressed as bars and SD.

A main effect for movement phase was observed (F_(1,156)_= 16.23, *p* < 0.001). Transporting the hand to the mouth showed a more positive RP than returning to the standby position (*p* < 0.001), indicating a preference of the forearm use in this phase (Figure 5B). Moreover, an interaction between movement phase and the moment of hand kinematics was also found (F_(2,156)_= 5.67, *p* < 0.01). *Post hoc* showed that during the transport phase there was a higher RP in the peak velocity of the hand as compared to the end of movement (*p* < 0.01), and to all moments of hand movement in the returning phase (beginning: *p* = 0.001, peak velocity: *p* < 0.001, ending: *p* < 0.01). Finally, during the transport phase the RP was also more positive in the beginning of movement compared to the peak velocity of the return (*p* < 0.05) (Figure 5B).

## 4 Discussion

We recorded the kinematics of bringing a plastic cup to the mouth and returning the upper limb to the side of the body at a standing position. The analysis showed that both the strategy for controlling hand motion and the pattern of coordination changed as a function of the phase of movement (transport or return). When the hand was moving towards the mouth, we observed that the TPV (hand acceleration time) and the TL were shorter compared with the return movement. Moreover, control subjects showed a common pattern of arm-forearm coordination throughout the transport movement while in the returning phase the pattern of movement differed among subjects. Those results reflect the effect of task goal constraints over the control of the hand movement and arm-forearm coordination.

In addition to the effects in movement phases, different RP patterns were expected to occur across groups. As the hand kinematics depends on the coordination between shoulder and elbow joints, we hypothesized that the muscle strength deficit caused by a TBPI would modify the arm-forearm coordination so as to conserve hand kinematics. In fact, we observed no differences between TBPI individuals and controls in relation to the PV, TPV and TL. In comparison to the control group, kinematic alterations in the injured UL were presented as a longer MD, more NMU and smaller NEH. Those changes point to an effort to accomplish the task with the injured UL. Kinematic analysis of coordination showed that the injured UL had a more negative RP when compared to control and uninjured UL movements, indicating that they might have used a different arm-forearm coordination strategy to move their hands. In addition to the changes in injured UL performance, differences in uninjured UL kinematics were observed. Despite no differences in the RP between the uninjured UL and control group were observed, for the uninjured UL a greater phase angle area was observed both for the arm and the forearm during the movement deceleration.

### 4.1 Goal Directed Movements: Task Goal Influences over UL Kinematic Performance

In the transport phase, the TPV was reduced in comparison to the return movement. This shorter value reflects a prolonged hand deceleration. The TPV parameter is known to reflect the motor system strategy adopted to control the movement (Marteniuk et al., 1987; Papaxanthis et al., 1998; Sartori et al., 2011). Shortening of the deceleration phase happens when the reaching movement has a specified target (de los Reyes-Guzmán et al., 2014). Marteniuk et al. (1987) showed that the presence of a target in a task creates a demand for precision in the control of actions. The more the reaching movements demand precise pointing or grasping, the longer is the deceleration phase of reaching (Marteniuk et al., 1987). Thus, in our task, the participant’s hand decelerates as the plastic cup comes closer to the mouth, so that it can match target specifications more precisely.

Previous studies have shown that the coordination between shoulder and elbow joints has a significant importance to the control of hand movements (Morasso, 1981; Soechting and Lacquaniti, 1981; Atkeson and Hollerbach, 1985). For each instant of UL motion, the motor system must control the motion of these joints in an orchestrated manner, regulating the rotations at the arm and at the forearm in a manner that results in a hand movement that attends task demands (Soechting and Lacquaniti, 1981). The use of the RP allows us to visualize how the motor system makes adaptations in motor plans when individual and environment constraints related to the task are presented (Barela et al., 2000; Daunoravičienė et al., 2017).

During the transport phase the RP was significantly more positive compared to the return, reflecting a greater forearm use in this movement phase. Moreover, the RP in the hand peak velocity was significantly more positive than the other moments (start and end) of hand movement. During the transport phase, the RP became progressively more positive as the hand accelerated. This positive RP in hand peak velocity indicated that the forearm had a greater relative contribution (in relation to the arm) to the generation of the hand peak velocity. This contribution is seen in Figure 4 (A and B). The RP started to decrease instants before the peak velocity, indicating that the forearm started to lose movement predominance when the deceleration phase approximated. As the hand came closer to the end of movement, the RP also decreased nearly to zero, suggesting that the arm displacement contributed to the final adjustments of the hand to the target.

This movement strategy is in line with evidence that the motor system plans hand movements in a way to generate a short straight trajectory towards the task target (Morasso, 1981; Soechting and Lacquaniti, 1981; Haggard et al., 1995). Indeed, we observed that the transport phase had a shorter TL in comparison to the return. In the proposed task if the participants had started the movement with the arm, a curved hand path with a longer trajectory would have been generated, instead of a shorter straight path. In their observations of UL reaching coordination, Haggard et al. (1995) suggested that the motor system would select a principal joint whose motions would cover most of the space between start and target positions, and then would use the other joints to produce hand-space regularities. This principle was observed in the kinematic analysis of UL multi-joint motion during daily living tasks (Dounskaia et al., 2020). Hand motion was led by the shoulder or the elbow depending both on the task and on the UL starting position, while other degrees of freedom were used to orient hand position (Dounskaia et al., 2020). In our analysis, the forearm led the acceleration of the transport phase, while the arm was used to the final adjustments of movement. This suggests that the presence of the target and the starting position of the limb constrained the coordination to a common pattern among participants.

Contrasting to the transport phase, the arm-forearm coordination strategies differed among participants during the UL return to the standby position (Figure 4D, 4E). The kinematic analysis of a drinking task in the work of Dounskaia et al. (2020) showed that participants’ strategy for the returning movement was to use the joints in the inverse order of the transport movement, which was not observed in our analysis. This difference might have arisen because in their work the returning movement had a specified endpoint, and this may have induced the selection of a particular coordination pattern. The returning movement in our task had no specific target and no constrictions, thus participants were free to choose the most convenient pattern of coordination according to their internal demands. Moreover, gravity has been shown to affect hand trajectory in downward motions (Papaxanthis et al., 1998), and also to facilitate downward UL control (Wang and Dounskaia, 2016).

### 4.2 Hand Kinematics of the Injured UL

In the present study participants were asked to reproduce the end position displayed in the figure in front of them in a comfortable self-selected speed, so as to allow their best possible performance. The injured UL movement presented a longer MD when compared to the control group. Temporal motor decisions follow a speed-accuracy trade off: movements of longer duration tend to have more spatial accuracy (Wolpert and Landy, 2012). The strength loss caused by TBPI can create difficulties in the control of UL movements, and as a consequence an adjustment in MD could facilitate better task accuracy. Although the task was performed more slowly for the injured UL group, a similar PV was achieved as compared to the control group. It is worth mentioning that the higher MD displayed by TBPI participants might reflect the need to make more corrections during movement execution with the injured UL, as seen by a higher NUM. The regular hand movement has a bell-shaped velocity curve with a single peak, and an increase in this number of peaks is interpreted as hand spatial corrections during ongoing movements (Kamper et al., 2002; de los Reyes-Guzmán et al., 2014; Bustrén et al., 2017). Beyond demanding more movement corrections, the muscle force deficit in the injured UL may also be the reason for the lower hand NEH. However, the mean NEH for injured UL was 89% of subjects’ height, which is a small difference to the performance of the control group (93%) and uninjured UL (94%). Taken together these results may reflect an effort of the motor system to preserve hand kinematics and achieve the task goal. This perspective is supported by the lack of any differences between groups for PV, TPV and TL.

In addition to the adjustments in hand kinematics, the TPV SD for the injured UL was five times higher than controls and three times higher than the uninjured UL in the transport phase. Probably, this higher SD may be a consequence of the clinical presentation of TPBI individuals, in which injured UL participants had to find a solution to control hand movements according to their injury severity and to their individual UL muscle strength.

In Souza et al. (2021), the uninjured UL kinematics was tested in a task involving trunk displacement. Results revealed altered kinematics for MD, PV, TPV and TL. TBPI individuals also showed lower TPV values in a bring-a-cup-to-the-mouth task, pointing to a more controlled hand deceleration (Souza et al., 2021). The difference observed in the previous study is justified by the injured UL variability in the present study. Further investigation is necessary to fully understand the effects of TBPI in uninjured UL kinematics during tasks that do not involve the trunk.

### 4.3 Alterations in the Coordination Pattern of TBPI Individuals

The injured UL showed a more negative RP in comparison to controls and to the uninjured UL independently of the movement phase. A more negative RP indicates that the injured UL movement relied more on the arm as opposed to the preferred forearm use found for the controls and the uninjured UL groups. However, careful analysis of individual RP curves revealed the existence of distinct arm-forearm coordination patterns among injured UL participants, and individual performances must be considered to the interpretation of the results. One participant in the injured UL group, which showed a very low score for shoulder muscles, showed a coordination pattern that relied mostly on arm movements. Three other injured UL participants had an RP curve that stayed below 20 degrees during the transport movement (Figure 4C), showing a coordination pattern that clearly used less forearm movements than controls and the uninjured UL group (Figure 4A, 4B). These changes in the injured UL arm-forearm coordination, associated with the effort of preserving hand function, are in accordance with results gathered in animal models showing that a peripheral nerve injury induces a joint kinematics reorganization that preserves the effector function (Chang et al., 2009, 2018; Sabatier et al., 2011; Bauman and Chang, 2013). Similarly, this principle of conserving effector function could explain our results. While the UL coordination displayed a reduction in forearm displacement in at least four of the injured UL participants, the preserved hand kinematics parameters PV, TPV and TL could be taken as resulting from an effort to conserve effector function.

The deficit in the shoulder muscle strength could be inducing this reduction in forearm displacements, because elbow motions generate an interaction torque at the shoulder joint that needs to be compensated by muscle contraction (Hollerbach and Flash, 1982; Ghez and Sainburg, 1995). Furthermore, it has been shown that these shoulder stabilizer muscles activate before the muscles that promote UL motion in reaching movements (Ricci et al., 2015). Maeda et al. (2020) showed that shoulder muscle activity can anticipatedly adapt when new constraints are introduced to the shoulder joint.

In deafferented patients, shoulder muscles presented a deficit in the ability to adapt muscle activation to those interaction torques (Ghez and Sainburg, 1995). TBPI may promote the learning of new intersegmental dynamics in daily tasks, because UL acquires a new configuration: muscle strength may not be fully recovered after surgery, and the deficit in proprioception could alter the feedback responses to interaction torques. It has been observed that TBPI individuals use more trunk motions when performing UL movements in different directions (Crouch et al., 2016; Webber et al., 2019; Nazarahari et al., 2020). Faity et al. (2021) showed that trunk motions occur to increase the reserve of anti-gravity shoulder torque. In fact, Crouch et al. (2016) observed that TBPI individuals have less reserve of shoulder strength in UL activities. So, instead of using the forearm to initiate the movement and raise interaction torques at the shoulder, the injured UL could be using the trunk and the arm to initiate movement while other joints assume movement stability and make the adjustments of hand position in space.

Alterations in the coordination pattern of TPBI individuals could also be a result of the plasticity induced by the injury (Mano et al., 1995; Iwase et al., 2001; Hsieh et al., 2002; Malessy et al., 2003; Pawela et al., 2010; Sokki et al., 2012; Yoshikawa et al., 2012; Liu et al., 2013; Qiu et al., 2014; Fraiman et al., 2016; Bhat et al., 2017). The reduced local connectivity in UL representations in M1 of TPBI individuals suggests a disturbance in the horizontal connections (Fraiman et al., 2016), pointed to have an important role in combining different joint movements (Huntley and Jones, 1991). This reduced activity in horizontal connections could go along with an alteration of upper limb movement synergies, where shoulder muscles would act in dissociation to the other joints during the movement.

During the return movement, the RP curves had a similar pattern to those observed in controls and in the uninjured UL, except for one individual whose RP curve was more negative than other injured UL participants (Figure 4C). The variability in the injured arm-forearm patterns reinforces that the UL coordination may be subjective to one’s internal demands when the task has no delimited endpoint.

Beyond injured UL arm-forearm altered coordination, the uninjured UL phase angle also presented curves that differed from controls. Although the phase angle curves of the injured UL displayed higher values through time (Figure 3 C,D,E,F), the areas measured under the curve did not differ statistically from controls and the uninjured UL groups. The absence of statistical difference in the injured group in comparison to the control and uninjured UL group could be due to the high variability presented in phase angle area. In the transport phase of the uninjured UL, it was seen that the forearm phase angle area was increased during deceleration. This result indicates that discrete alterations in forearm kinematics may be present in this limb, but not to the extent of causing significant alterations in arm-forearm coordination. In our previous work it was seen that uninjured UL movements have an increased demand of precision when trunk displacement was involved (Souza et al., 2021). During the return phase of the present experiment, both the arm and forearm phase angles areas of the uninjured UL were increased when compared to controls. Since the returning movement had no target, these alterations might reflect a movement adapted to new internal demands, for example the changes in balance observed in TBPI individuals (Souza et al., 2016). TBPI individuals display alterations in UL motor representations of both hemispheres (Liu et al., 2013; Fraiman et al., 2016; Torres et al., 2018; Rangel et al., 2021). Recent findings suggest that the neural activity in the motor cortex of single hemisphere allows decoding the kinematics from both the contralateral limb and the ipsilateral limb, suggesting that movements may be bi-hemispherically represented in humans (Bundy et al., 2018). Besides, in stroke patients movement training of the ipsilesional limb improved the kinematic performance of the contralesional limb (Pohl and Winstein, 1999; Maenza et al., 2021).

### 4.4 Considerations for Rehabilitation

The analysis of injured UL kinematics after a TBPI demonstrated that the natural pattern of arm-forearm coordination was disturbed. Previous studies have shown that shoulder muscles are important to the stability of UL movements (Ricci et al., 2015; Maeda et al., 2020). We suggest that the lack of shoulder strength in the injured UL may be the reason for the changes in arm-forearm coordination. Usually, therapeutic approaches for restoring TPBI UL function aims to improve elbow and shoulder muscle strength by restorative surgeries and physical therapy, with a higher priority given to biceps recovery. Gaining shoulder muscle strength might be important to reduce the compensatory movements observed in previous studies (Webber et al., 2019; Nazarahari et al., 2020). Moreover, the alterations in the kinematics of the uninjured UL gives evidence for the inclusion of the uninjured UL in rehabilitation programs. The idea of a broad rehabilitation approach has already been suggested by previous studies (Souza et al., 2016, 2021), which pointed to a rehabilitation program that included balance assessment and considered the uninjured UL for training

### 4.5 Study Limitations

The TBPI individuals presented heterogeneity regarding the extent of the injury, the type of surgical procedure performed, time elapsed from the lesion, and muscle strength at the time of experiment. Only a small sample of participants was able to perform movements with the injured UL, and this taken together with the high variability in their data may have hidden some previous findings in the uninjured UL (Souza et al., 2021). Future analysis may be improved with a higher number of participants able to perform injured UL movements and by the categorization of the severity of the injury in kinematic analysis. Also, the analysis of trunk motions would help identify the adaptations in the coordination of injured UL movements.

## 5 Conclusion

Our results indicate that the arm-forearm coordination in the injured UL is changed in order to preserve hand function. Although the patterns of coordination were adjusted to individual’s injury severity and UL strength, most of the injured UL performances pointed to a reduction in the forearm contributions to the movement, which probably occurred because of a reduction in shoulder strength. These results highlight the importance of restoring shoulder function by surgeries and rehabilitation. Finally, the kinematics of the uninjured UL seems also to be affected after a TBPI, as suggested by the arm and forearm motion departure from that of controls.

## 6 Acknowledgements

The authors would like to thank Gabriel Freire Miranda, Lidane Souza, José Vicente Martins and José Magalhães de Oliveira for their contribution to this work. This research was supported by the Conselho Nacional de Desenvolvimento Científico e Tecnológico, CNPq (grant number 310397/2021), Fundação de Amparo à Pesquisa do Estado do Rio de Janeiro FAPERJ (grants E26/010002474/2016, CNE 202.785/2018 and E-26/010.002418/2019), and Financiadora de Estudos e projetos FINEP (PROINFRA HOSPITALAR grant 18.569-8). This research is also part of the activities of the Fundação de Amparo à Pesquisa do Estado de São Paulo FAPESP’s Research, Innovation and Dissemination Center for Neuromathematics-NeuroMat (FAPESP grant 2013/07699-0). Luiggi Lustosa is a PhD scholarship holder from Conselho Nacional de Desenvolvimento Científico e Tecnológico, CNPq (process number: 160457/2018-1).

