## Supplementary tables for "Upper limb joint coordination acts to preserve hand kinematics after a traumatic brachial plexus injury"

### Supplementary material

**Table S1.** Hand kinematic performance

| | **Movement Duration (mean ± SD, s)** | | |
|---|---|---|---|
| | Control | Uninjured UL | Injured UL[(a)] |
| Transport | 1.07 ± 0.23 | 1.20 ± 0.24 | 1.47 ± 0.47 |
| Return | 1.28 ± 0.30 | 1.43 ± 0.24 | 1.43 ± 0.27 |
| | **Peak Velocity (mean ± SD, mm/s)** | | |
| | Control | Uninjured UL | Injured UL |
| Transport | 2230.34 ± 516.39 | 1945.65 ± 443,89 | 1779.17 ± 276.73 |
| Return | 1907.76 ± 458.99 | 1595.42 ± 415.21 | 1799.91 ± 540.32 |
| | **Time to peak velocity – TPV (mean ± SD)** | | |
| | Control | Uninjured UL | Injured UL |
| Transport[(b)] | 0.37 ± 0.02 | 0.34 ± 0.03 | 0.37 ± 0.10 |
| Return | 0.43 ± 0.06 | 0.41 ± 0.06 | 0.39 ± 0.13 |
| | **Trajectory length – TL (mean ± SD, mm)** | | |
| | Control | Uninjured UL | Injured UL |
| Transport[(c)] | 1154.51 ± 78.11 | 1101.75 ± 109.90 | 1115.51 ± 120.64 |
| Return | 1289.58 ± 107.14 | 1197.19 ± 143.27 | 1227.62 ± 179.47 |
| | **Normalized end height – NEH (mean ± SD)** | | |
| | Control | Uninjured UL | Injured UL[(d)] |
| Transport | 0.93 ± 0.02 | 0.94 ± 0.02 | 0.89 ± 0.04 |

[(a)] this group was significantly different from controls ($p < 0.05$)
[(b)] the transport phase was significantly different from the return phase ($p < 0.01$)
[(c)] the transport phase was significantly different from the return phase ($p < 0.001$)
[(d)] Injured UL was significantly different from uninjured UL and controls ($p < 0.05$)

**Table S2.** Averages of RP time windows

**Transport to mouth**

| **Hand moment** | **Relative phase (mean ± SD, deg.)** | | |
|---|---|---|---|
| | Control | Uninjured UL | Injured UL[a] |
| Start[b] | 11.62 ± 3.51 | 12.90 ± 3.93 | 4.18 ± 11.37 |
| Peak[c] | 28.52 ± 7.64 | 31.66 ± 8.34 | -0.40 ± 67.71 |
| End | -0.51 ± 2.82 | 2.04 ± 3.72 | -10.77 ± 34.34 |

**Return to standby**

| **Hand moment** | **Relative phase (mean ± SD, deg.)** | | |
|---|---|---|---|
| | Control | Uninjured UL | Injured UL[a] |
| Start | 3.18 ± 4.04 | -1.17 ± 3.10 | -14.49 ± 34.78 |
| Peak | 1.31 ± 27.33 | -9.48 ± 20.26 | -14.44 ± 51.11 |
| End | 2.62 ± 3.88 | 2.48 ± 3.43 | -13.24 ± 38.90 |

[a] significantly different from controls ($p < 0.001$) and uninjured UL ($p < 0.001$)
[b] transport start differed significantly from return peak velocity ($p < 0.01$)
[c] transport peak velocity differed significantly from transport end ($p < 0.01$), return start ($p = 0.001$), return peak velocity ($p < 0.001$), and return end ($p < 0.01$).
